# *Nosema ceranae* infection disrupts nestmate recognition in honey bees

**DOI:** 10.64898/2026.08.02.742297

**Authors:** Yujie Li, Honghong Li, Xusheng Li, Zachary Y Huang

## Abstract

Social immunity reduces pathogen transmission in eusocial insects by collectively detecting and excluding infected or foreign individuals. In honey bees (*Apis mellifera*), guard workers enforce colony boundaries through nestmate recognition. Here we show that infection with the microsporidian parasite *Nosema ceranae* weakens this defense. Across 600 behavioral assays, infected workers were 45% less likely to be rejected than uninfected controls. Infection was also associated with modest but significant shifts in cuticular hydrocarbon profiles, the primary cues underlying nestmate recognition. Multivariate analyses revealed modest but significant treatment-associated differences in CHC composition that were not attributable to dispersion effects. These findings demonstrate that infection can degrade the reliability of recognition cues and thereby compromise a core behavioral component of social immunity.

## INTRODUCTION

Group living imposes substantial epidemiological costs: dense aggregations and frequent contact increase opportunities for parasite transmission. Eusocial insects have therefore evolved “social immunity” -- collective defenses that limit pathogen establishment and spread through coordinated behaviors and organization (Schmid-Hempel 1998; Cremer et al. 2007; Wilson-Rich et al. 2009; Meunier 2015). Although much work has focused on within-nest processes such as grooming and spatial reorganization (Ugelvig & Cremer 2007; Stroeymeyt et al. 2018), disease risk is also shaped by movement between colonies. In honey bees (*Apis mellifera*), drifting and robbing can transport parasites among neighboring hives, making colony boundary defense a critical component of social immunity (Breed et al. 2004b; Pfeiffer & Crailsheim 1998).

Guards enforce colony boundaries by inspecting arriving workers and attacking individuals perceived as non-nestmates. Recognition is typically framed as template-based signal detection, in which an encountered chemical profile is compared against an internal representation of colony odor and aggression is expressed when similarity falls below a threshold (Reeve 1989; Sherman et al. 1997; van Zweden & d’Ettorre 2010). Recognition accuracy therefore depends on reliable signal production and sufficient separation between nestmate and non-nestmate cue distributions.

In social insects, cuticular hydrocarbons (CHCs) constitute the primary recognition cues (Howard & Blomquist 2005; van Zweden & d’Ettorre 2010; Leonhardt et al. 2016). These compounds are products of lipid metabolism and vary with age, task and physiological condition (Blomquist & Bagnères 2010; Kather et al. 2011). Because guards rely on quantitative similarity assessment, even modest shifts in CHC composition can alter acceptance decisions (Dani et al. 2001). Processes that perturb metabolic state may therefore reduce cue reliability and increase recognition error.

Pathogens are strong candidates to generate such perturbations. Infection can disrupt host physiology, alter behavior and modify chemical communication (Alaux et al. 2012; Geffre et al. 2020). Whether infection compromises colony boundary defense by altering host cues, guard decision rules, or both, remains insufficiently tested.

The microsporidian *Nosema ceranae* infects the midgut epithelium of adult honey bees (Higes et al. 2006; Fries 2010) and induces energetic stress and behavioral changes (Mayack & Naug 2009; Goblirsch et al. 2013). Because CHC biosynthesis is metabolically grounded, such physiological disruption could shift recognition cues. If infection reduces chemical distinctness, guards may accept infected non-nestmates more frequently, weakening this first line of defense against intercolony transmission.

Here we test whether infection reveals a vulnerability in colony boundary defense. Using controlled infections and standardized guard assays, we quantify how *N. ceranae* affects rejection of non-nestmates and assess infection-associated shifts in CHC composition. We predict that infected non-nestmates will be rejected less often than controls and that infection will be associated with measurable changes in chemical profiles, linking parasite-induced physiological perturbation to degraded recognition accuracy.

## MATERIALS AND METHODS

### Study colonies and experimental design

Experiments were conducted during the active beekeeping season at the Michigan State University apiary (East Lansing, MI, USA) during summer of 2015. Colonies were maintained in standard Langstroth hives and managed according to standard apicultural practices. Colonies used in behavioral assays were queenright, contained brood at all stages, and exhibited normal foraging activity at the time of testing. Source colonies providing experimental bees were treated with fumagillin 3 months prior to experiment, using manufacturer recommended doses.

To generate age-controlled experimental bees, brood frames containing late-stage pupae were removed from four source colonies and placed in an incubator at 34.5 °C and 50% relative humidity. Newly emerged workers (<24 h old) were collected and randomly assigned to infection or control treatments (110 bees per cage). Bees from each source colony were maintained separately by treatment in ventilated cages supplied ad libitum with 50% (w/v) sucrose solution and water. Cages were held in darkness at 34.5 °C.

### Spore preparation and experimental infection

*Nosema ceranae* spores were obtained from naturally infected donor colonies maintained separately from experimental colonies. Midguts were dissected from infected workers in distilled water and homogenized using a sterile glass tissue grinder. The homogenate was filtered through sterile gauze to remove debris and centrifuged at 10,000g for 2 min. The pellet was resuspended in distilled water and spore concentration determined using a hemocytometer under phase-contrast microscopy at 400× magnification.

Spore suspensions were adjusted to 25,000 spores µL^-1^. Each newly emerged worker assigned to the infection treatment received 2 µL of spore suspension (50,000 spores per bee) delivered individually using a calibrated micropipette. Bees were starved for 2 h prior to feeding and consumed the droplet voluntarily. Control bees received 2 µL of sterile 50% sucrose solution without spores. Bees were placed individually in glass tubes for 30 min to prevent trophallaxis that could alter the inoculation dose. After that, all infected bees were moved to one cage while all control bees were moved to another cage. The two cages were placed in an incubator at 34.5°C and 50% RH.

Bees were maintained for seven days post-inoculation to allow infection to develop. Infection status was verified in a subsample of 30 treated and 30 control bees by dissecting midguts and quantifying spores microscopically using a hemocytometer. Infection prevalence exceeded 93% in treated bees and no spores were detected in the control bees.

### Guard aggression assays

Nestmate recognition was quantified using a paired (two-bee) laboratory bioassay following Breed et al. (2004a). Guard bees were collected directly from hive entrances during peak foraging hours (09:00–14:00) and immediately transferred to 5 mL clear glass vials. Guards were identified by their location at the hive entrance and characteristic inspection behavior toward incoming bees.

Each guard was paired with a single 8-day-old introduced worker (infected or control) from a non-nestmate source colony. Introduced bees were briefly chilled on ice (<60 s) to facilitate handling and then placed into the vial containing the guard. Interactions were observed for 5 min.

Aggression was scored as a binary response: aggressive (stinging attempt or sustained biting) or non-aggressive (inspection without attack, antennation, or no interaction), within a 5 min observation window. Observations were recorded in real time by an observer blind to the treatments.

For each focal guard bee colony, 25 infected and 25 control non-nestmates from each source colony were tested, with a total of three guard colonies tested, yielding a total of 600 assays (25 bees x 3 guard colonies x 2 treatments x 4 source colonies = 600, Fig. 1). Trials were randomized with respect to treatment order and source colony to prevent temporal bias.

**Fig. 1.**
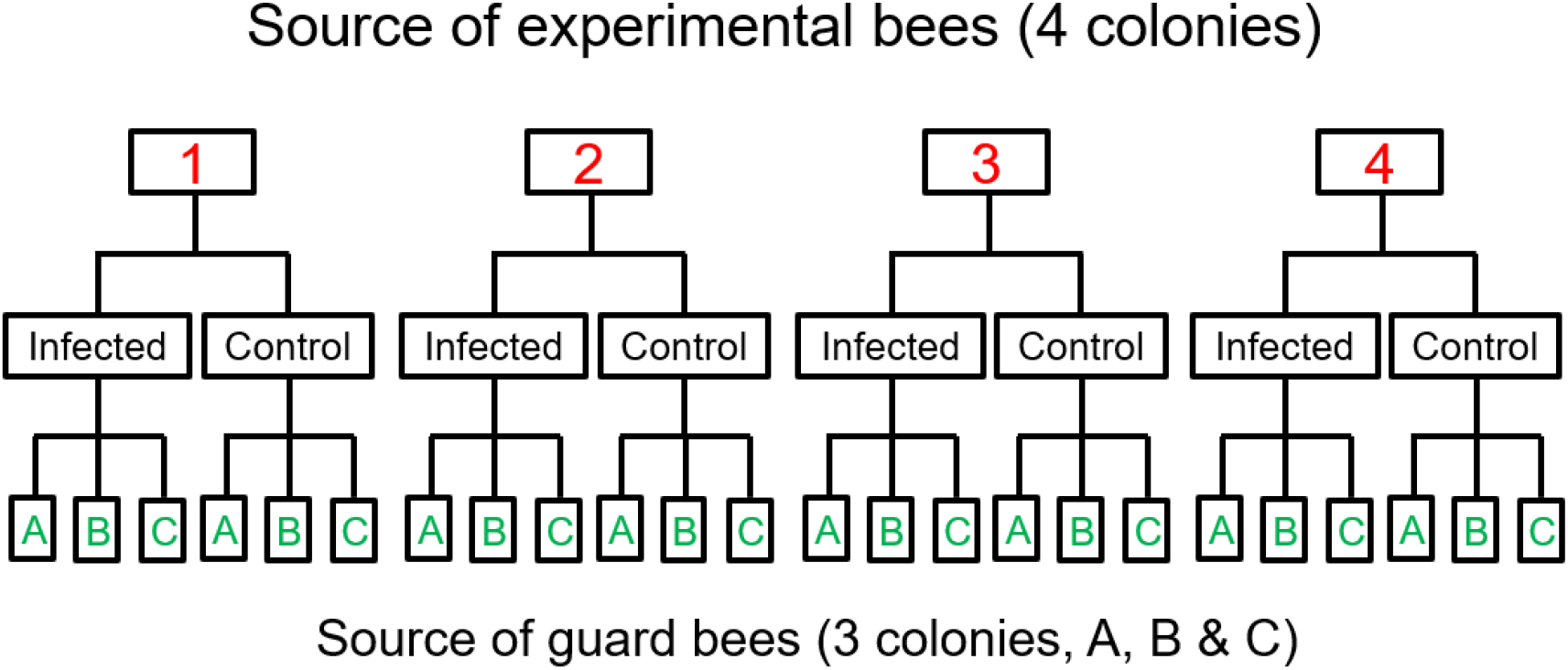
Experimental design showing four source colonies providing infected and control workers (both 8 days old) that were tested against guards from three unrelated colonies.

### Cuticular hydrocarbon extraction

Cuticular hydrocarbon analyses were conducted in 2018 in Guangxi, China using workers from two additional colonies (colonies 5 and 6), independent of those used in the behavioral assays. Bees were infected using the same dosage of *Nosema ceranae* as the guard bee experiment, and the infected and control bees, both 8 day old, were freeze-killed at −20 °C and allowed to equilibrate to room temperature prior to extraction.

Each individual bee was immersed in 2 mL of HPLC-grade hexane (≥99% purity, Chengdu Kelong Chemical Co. Ltd, Chengdu, China) and vortexed for 2 min. Each bee was then removed and solvent extracts were transferred to a clean glass vial.

Extracts were evaporated under a gentle stream of nitrogen gas to near dryness and reconstituted in 200 µL of hexane. Samples were stored at −20 °C in sealed vials with PTFE-lined caps until GC/MS analysis.

### GC/MS analysis

CHC extracts were analyzed using an Agilent 7890A-5975C gas chromatography–mass spectrometry (GC/MS) with an Agilent 122-5532UIDB-5MS column (30 m×250 µm×0.25 µm) and EI ion source. The injection volume was 1 µL. The GC parameters were set as the following: injection temperature 250 ºC, column temperature 80 °C, hold for 3 min, then increase at a rate of 10 °C to 250 °C, hold for 20 min, sampling time 60 min, gas flow rate 1 mL/min. For MS, the following parameters were used: full scan mode, ion scan range 50-600, ion source temperature 230 ºC, MS quadrapole temperature 150 ºC.

Between every 10 samples, 10 blanks were injected to monitor contamination and instrument drift. Internal standards were not used in this study, as the inclusion of standards would increase the difficulties of compound separation by GC and is disadvantageous for identification of CHC in samples.

Chromatographic peaks were detected and integrated using MSD Productivity ChemStation (G1701EA; Agilent Technologies, Inc., Santa Clara, USA). Peaks were identified by comparison of mass spectra to NIST library databases and by reference to retention indices where available. Peaks were assigned to the same compound if their differences in absolute retention time (t_R_) are no more than 5% under the same chromatographic conditions.

### Statistical analysis of behavioral data

Guard rejection behavior was analyzed using generalized linear mixed-effects models with a binomial error distribution and logit link (function glmer, package lme4).

Aggressive responses (bite, sting attempt or sustained attack) were coded as 1 and non-aggressive encounters as 0. Infection status of the introduced worker (infected vs. control) was included as a fixed effect. Initial models included both source colony and guard colony as random intercepts. Because the estimated variance for guard colony was zero, the final model retained source colony as the sole random intercept. Analyses were conducted in R (version 4.5.2).

### Data processing of CHC data

Before GC/MS data processing, samples were screened for contamination from column coating (silicon compounds), and contaminated samples were excluded. Chromatographic peaks were retained only if they met predefined criteria for reproducibility, chromatographic separation, and signal quality (retention time deviation ≤ 5%, relative standard deviation ≤ 10%, resolution > 1.0, signal-to-noise ratio > 5), with silicon contaminant peaks removed. Peak abundances were converted to relative percentages of the total CHC signal for each individual, such that each sample summed to 100%. Two samples with abnormal total integrated peak abundance were excluded prior to normalization.

Analyses were conducted in R (version 4.5.2). Chemical profiles were analysed using Bray–Curtis dissimilarities calculated from relative peak abundances (function vegdist, package vegan). Patterns of multivariate variation were visualized using principal coordinates analysis (PCoA; function pcoa, package ape). Treatment effects on CHC composition were tested using permutational multivariate analysis of variance (PERMANOVA; function adonis2, package vegan) with 9,999 permutations. Because colony identity strongly influences hydrocarbon profiles, permutations were constrained within colony. Homogeneity of multivariate dispersion was assessed using betadisper with permutation tests (permutest, package vegan) to verify that significant PERMANOVA results reflected differences in group centroids rather than differences in within-group dispersion.

## Results

Infected workers were significantly less likely to be rejected than controls after accounting for variation among source colonies (β = -0.60 ± 0.19 SE, z = -3.21, P = 0.0013), corresponding to a 45% reduction in the odds of rejection (odds ratio, exp(β) = 0.55; Fig. 2).

**Fig. 2.**
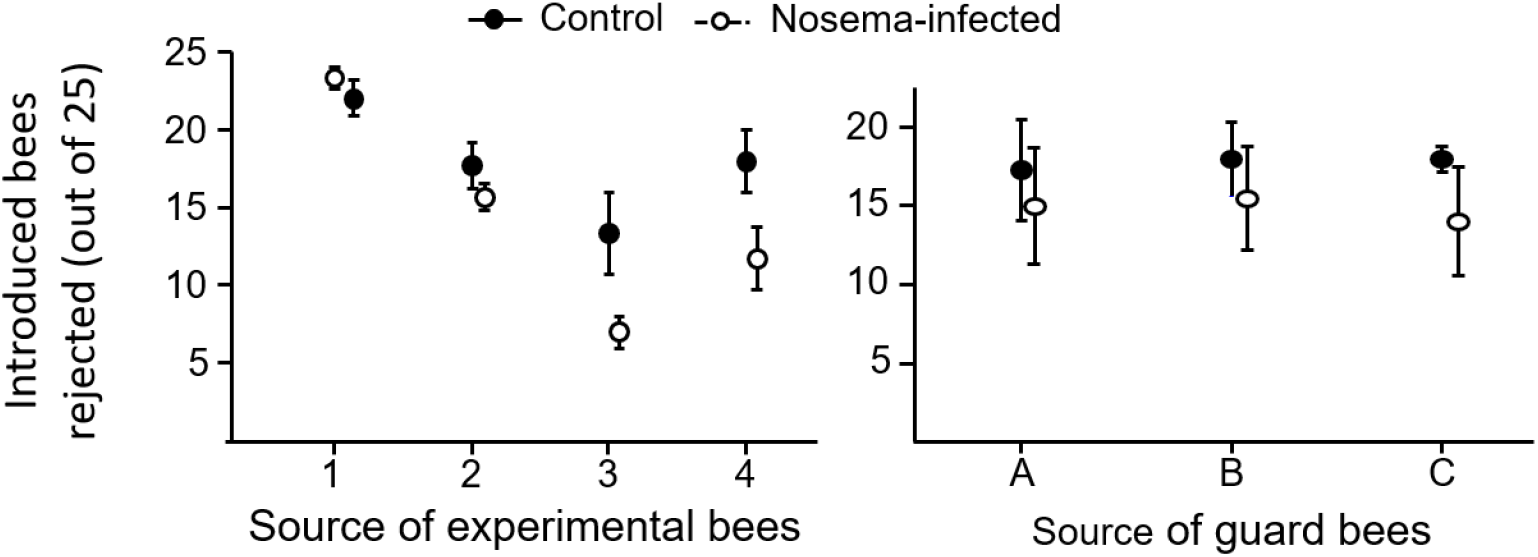
*Nosema*-infected workers (open circles) were rejected less frequently (means ± SE) than age-matched control workers (filled circles). Upper panel shows data grouped by source colony (n = 3 independent replicates per point); lower panel shows the same data grouped by guard colony (n = 4 independent replicates per point).

Infection was associated with a significant shift in CHC composition (PERMANOVA: R^2^ = 0.046, F = 3.99, P = 0.013; Fig. 3). Multivariate dispersion did not differ between treatments (PERMDISP, P = 0.105), indicating that the effect reflects differences in centroid location rather than variance heterogeneity. The first two PCoA axes accounted for 55.0% and 20.3% of the variation, respectively.

**Fig. 3.**
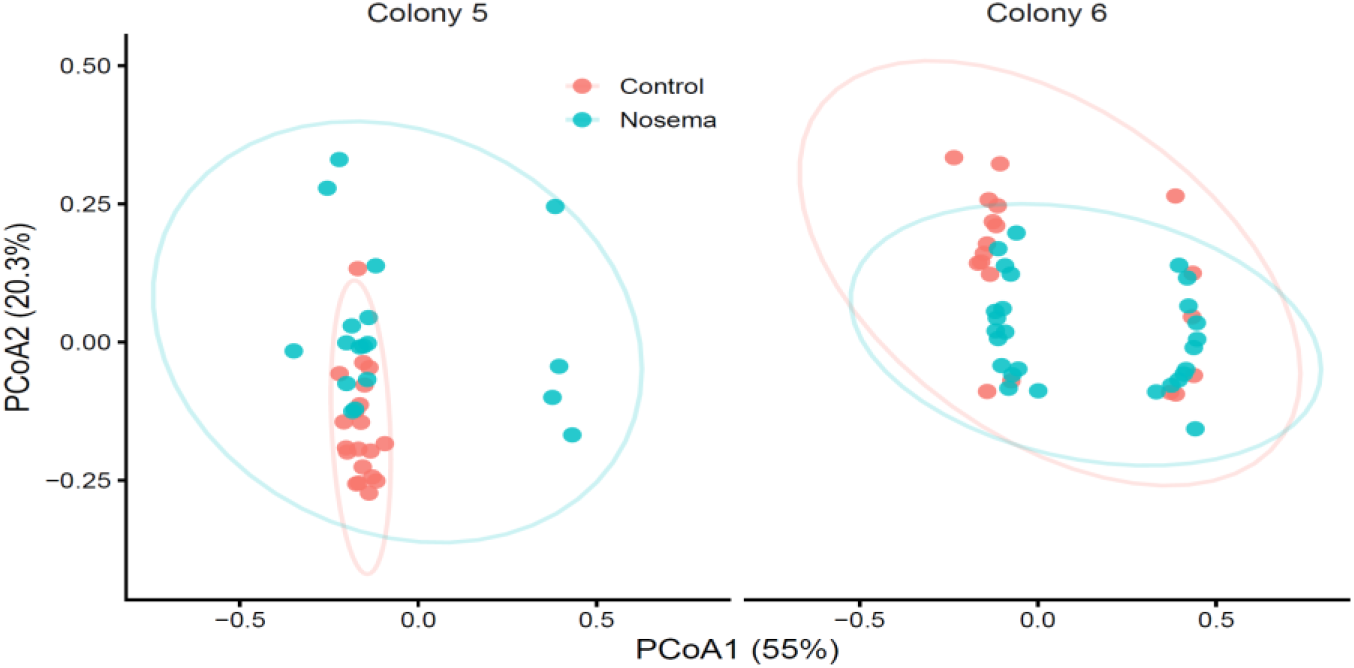
Principal coordinates analysis (PCoA) of cuticular hydrocarbon (CHC) profiles. Infection with *Nosema* was associated with a significant shift in CHC composition (PERMANOVA, P = 0.013), while colony identity remained an important source of variation. Ellipses represent 95% confidence regions.

## Discussion

Our results show that *Nosema ceranae* infection weakens colony boundary defense by increasing acceptance of infected non-nestmates. Because nestmate recognition is a core component of social immunity, this finding identifies a vulnerability at the colony’s epidemiological interface. Infection was associated with modest but significant shifts in cuticular hydrocarbon (CHC) composition -- the primary cues mediating recognition in social insects -- supporting the interpretation that pathogen-induced physiological perturbation can degrade cue reliability and increase recognition error.

From a signal-detection perspective, template-based recognition depends on stable cue distributions and appropriately calibrated decision thresholds (Reeve 1989; Sherman et al. 1997; van Zweden & d’Ettorre 2010). Even small coordinated shifts in hydrocarbon composition can alter similarity relationships in a multidimensional cue space, increasing overlap between nestmate and non-nestmate distributions. Our results therefore suggest that such recognition systems may be intrinsically sensitive to physiological disturbance.

This sensitivity has epidemiological consequences. Social immunity is often framed in terms of within-nest processes such as grooming and spatial reorganization (Cremer et al. 2007; Stroeymeyt et al. 2018), but effective defense also depends on the integrity of colony borders. In apiaries where worker drifting is common (Pfeiffer & Crailsheim 1998), even modest increases in acceptance of infected foreign bees could facilitate between-colony transmission.

CHC production is rooted in lipid metabolism and reflects physiological state (Howard & Blomquist 2005; Blomquist & Bagnères 2010). Because *N. ceranae* disrupts nutrient absorption and induces energetic stress (Mayack & Naug 2009; Fries 2010), infection-associated shifts in hydrocarbons may arise as downstream consequences of metabolic perturbation. Elevated juvenile hormone titers in infected workers (Goblirsch et al. 2013; Lin et al., submitted) provide a plausible endocrine pathway linking infection to altered hydrocarbon biosynthesis. Whether such changes represent incidental physiological spillover or a form of host manipulation remains unresolved. Importantly, adaptive manipulation is not required for epidemiological impact: any infection-induced degradation of cue reliability can increase false-negative acceptance at the colony boundary. Similar pathogen-associated changes in recognition have been reported for honey bees following Israeli acute paralysis virus infection (Geffre et al., 2020). Together, these studies suggest that pathogen-induced disruption of nestmate recognition may represent a general consequence of infection-induced physiological perturbation rather than an idiosyncratic effect of a particular parasite.

More broadly, our findings highlight a general principle: communication systems that underpin collective defense are only as robust as the physiological processes that generate their signals. When those processes are perturbed by infection, the informational foundation of social immunity may erode. Because CHC-mediated recognition is widespread across eusocial insects (Howard & Blomquist 2005; van Zweden & d’Ettorre 2010; Leonhardt et al. 2016), similar vulnerabilities may occur wherever group membership is enforced through metabolically grounded chemical cues.

By linking infection, chemical signaling and boundary defense (Fig. 4), our study suggests that pathogens may influence disease dynamics not only through direct effects on host survival or contact structure, but also by degrading the reliability of the signals that regulate collective defense.

**Fig. 4.**
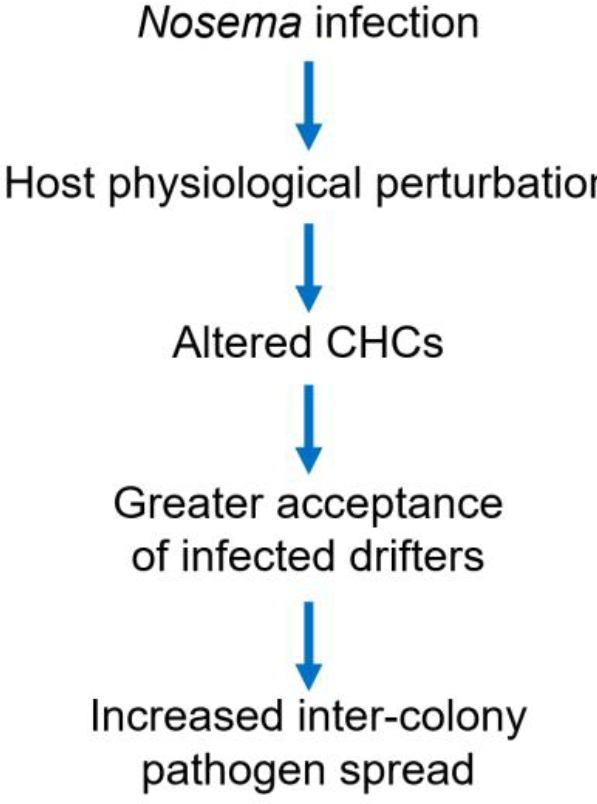
Conceptual model illustrating how *Nosema* infection may facilitate pathogen spread by altering cuticular hydrocarbon cues and increasing acceptance of infected non-nestmates by guard bees.

## Code availability

Data used to generate Figures and R code for statistical analyses and figures are available at: https://doi.org/10.5281/zenodo.21399303

## Acknowledgements

We gratefully thank Jindong Zhang, Xianbing Xie and Qing Wang who provided help in the experiment. We also acknowledge the sources of funding: the National Natural Science Foundation of China (U2571211; 32270551; 32470541 to YL). Michigan Beekeepers Association provided generous support to ZYH.

## Author Contributions

YL: behavioral data acquisition, writing first draft, HL: *Nosema* infection for CHC experiment, SL: CHC data acqusition, ZYH: Conception, experimental design, data analysis, visualisation, editing and supervision, . All authors participated in writing/editing of the manuscript.

## References

Alaux, C., Le Conte, Y., Adams, H.A., Rodriguez-Zas, S., Grozinger, C.M. & Sinha, S. (2012). Effects of immunostimulation on social behavior, chemical communication and genome-wide gene expression in honey bee workers. BMC Genomics, 13, 558.

Blomquist, G.J. & Bagnères, A.-G. (eds.) (2010). Insect Hydrocarbons: Biology, Biochemistry, and Chemical Ecology. Cambridge University Press.

Breed, M.D., Diaz, P.H., Lucero, K.D. (2004a). Olfactory information processing in honeybee, Apis mellifera, nestmate recognition. Animal Behaviour, 68, 921–928.

Breed, M.D., Guzmán-Novoa, E. & Hunt, G.J. (2004b). Defensive behavior of honey bees: organization, genetics, and comparisons with other bees. Annual Review of Entomology, 49, 271–298.

Cremer, S., Armitage, S.A.O. & Schmid-Hempel, P. (2007). Social immunity. Current Biology, 17, R693–R702.

Dani, F.R., Jones, G.R., Destri, S., Spencer, S.H. & Turillazzi, S. (2001). Deciphering the recognition signature within the cuticular chemical profile of paper wasps. Animal Behaviour, 62, 165–171.

Fries, I. (2010). Nosema ceranae in European honey bees (Apis mellifera). Journal of Invertebrate Pathology, 103, S73–S79.

Geffre, A.C., Gernat, T., Harwood, G.P., Jones, B.M., Morselli Gysi, D., Hamilton, A. R., Bonning, B.C., Toth, A.L., Robinson, G.E., Dolezal, A.G. (2020). Honey bee virus causes context-dependent changes in host social behavior. Proceedings of the National Academy of Sciences of the United States of America, 117(19), 10406–10413.

Goblirsch, M., Huang, Z.Y., Spivak, M. (2013). Physiological and behavioral changes in honey bees (Apis mellifera) induced by Nosema ceranae infection. PloS one, 8(3), e58165.

Higes, M., Martín-Hernández, R. & Meana, A. (2006). Nosema ceranae, a new microsporidian parasite in honey bees. Journal of Invertebrate Pathology, 92, 93–95.

Howard, R.W. & Blomquist, G.J. (2005). Ecological, behavioral, and biochemical aspects of insect hydrocarbons. Annual Review of Entomology, 50, 371–393.

Kather, R., Drijfhout, F.P. & Martin, S.J. (2011). Task group differences in cuticular lipids in the honey bee Apis mellifera. Journal of Chemical Ecology, 37, 205–212.

Leonhardt, S.D., Menzel, F., Nehring, V. & Schmitt, T. (2016). Ecology and Evolution of Communication in Social Insects. Cell, 164, 1277–1287.

Lin, R., Sullivan, J., Webster, T.C., Huang, Z.Y. (Submitted). Altered juvenile hormone dynamics mediate earlier foraging in Nosema-infected honey bees.

Mayack, C. & Naug, D. (2009). Energetic stress in the honeybee Apis mellifera from Nosema ceranae infection. Journal of Invertebrate Pathology, 100, 185–188.

Meunier, J. (2015). Social immunity and the evolution of group living in insects. Philosophical Transactions of the Royal Society B, 370, 20140102.

Pfeiffer, K.J. & Crailsheim, K. (1998). Drifting of honeybees. Insectes Sociaux, 45, 151–167.

Reeve, H.K. (1989). The evolution of conspecific acceptance thresholds. The American Naturalist, 133, 407–435.

Schmid-Hempel, P. (1998). Parasites in Social Insects. Princeton University Press.

Sherman, P.W., Reeve, H.K. & Pfennig, D.W. (1997). Recognition systems. In: Behavioral Ecology: An Evolutionary Approach (eds Krebs, J.R. & Davies, N.B.). Blackwell Science.

Stroeymeyt, N. et al. (2018). Social network plasticity decreases disease transmission in a eusocial insect. Science, 362, 941–945.

Ugelvig, L.V. & Cremer, S. (2007). Social prophylaxis: group interaction promotes collective immunity in ant colonies. Current Biology, 17, 1967–1971.

van Zweden, J.S. & d’Ettorre, P. (2010). Nestmate recognition in social insects and the role of hydrocarbons. Insectes Sociaux, 57, 1–14.

Wilson-Rich, N., Spivak, M., Fefferman, N.H. & Starks, P.T. (2009). Genetic, individual and group facilitation of disease resistance in insect societies. Annual Review of Entomology, 54, 405–423.

